# Priors for perceived speed combine supramodally for visual-tactile motion but not for audio-visual motion

**DOI:** 10.64898/2026.09.17.751718

**Authors:** Alessia Tonelli, Cameron K. Phan, David Alais

## Abstract

Perception combines incoming sensory evidence with priors learned from preceding input. One example is the central tendency effect in which stimulus estimates regress towards a prior based on the mean of the stimulus distribution. Whether such priors are specific to each modality or shared across modalities remains unresolved. Here, we examined this question using speed perception in the visual, auditory and tactile modalities and used central tendency as a behavioral probe. By giving each modality its own speed range, clear predictions emerge when modalities are interleaved: if priors are modality specific, each modality should maintain its own central-tendency bias whereas a supramodal prior would exhibit a different mean based on the pooled set of speeds. Moreover, a supra-modal prior should exhibit precision weighting, with the more reliable cue receiving more weight. For vision and touch, speed estimates converged when interleaved: both modalities shifted but the less reliable one shifted more, consistent with a precision-weighted supramodal prior. For vision and audition, the priors remained largely distinct: audition (the less reliable modality) showed a partial shift but did not converge on the precision-weighted, supramodal mean. This was not explained by the large audio–visual reliability difference: it persisted when visual precision was reduced. These results show that in the motion domain, visual and tactile signals can combine crossmodally, but auditory and visual signals do not. Supramodal processing of motion therefore depends on which senses are involved and is likely explained by the lack of an early spatial mapping in audition.

## Introduction

Perception is the result of an inferential process in which the brain combines incoming sensory information with prior knowledge, allowing it to overcome the noise and ambiguity inherent in sensory inputs and build stable representations of the environment (Berniker et al., 2010; Knill & Pouget, 2004). Perception is therefore not fixed but context-dependent, often showing systematic biases that reflect prior experience. A well-known example is the central tendency effect (see Figure 1D): when observers estimate a physical attribute of a stimulus, their responses regress toward the mean of the presented stimulus range, overestimating low values and underestimating high ones (Hollingworth, 1910). The effect is observed across many perceptual domains (Corbin et al., 2017; Jazayeri & Shadlen, 2010; Olkkonen et al., 2014) and sensory modalities (Riskey et al., 1979; Tonelli, Mazzola, et al., 2025; Zimmermann & Cicchini, 2020), indicating that perception is sensitive to the global context in which a stimulus occurs and not specific to the stimulus itself.

**Figure 1.**
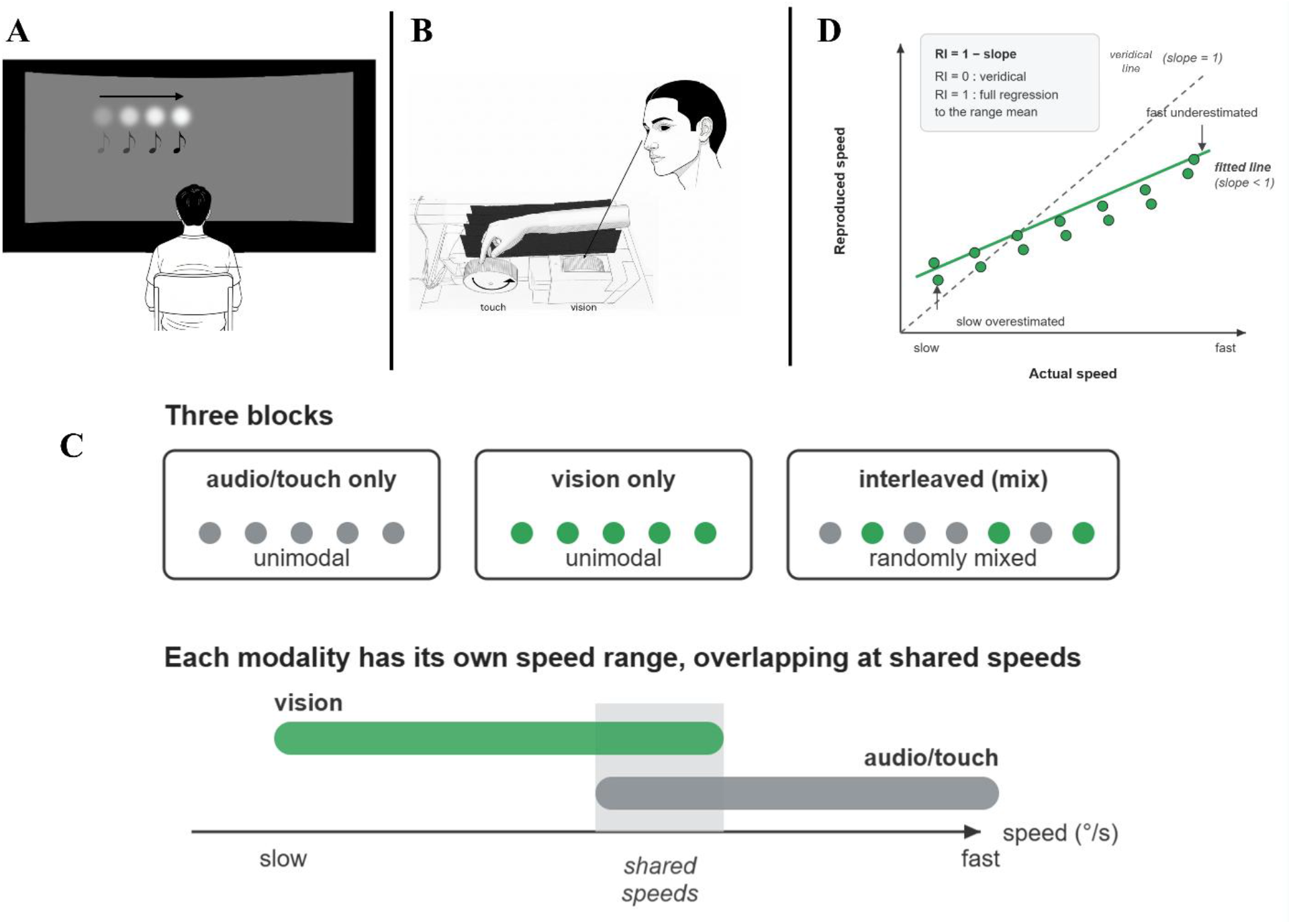
Apparatus and experimental paradigm. **(A)** Audio–visual set-up: participants viewed a disc moving horizontally across a projection screen (vision) or heard equivalent motion rendered over a horizontal speaker array (audition) and reproduced the perceived speed on a response bar. **(B)** Visual-tactile set-up: visual and tactile motion were delivered by two rotating wheels, the tactile wheel felt with the fingertip and the visual wheel viewed directly; speed was reproduced on the same response bar. **(C)** Paradigm (illustrated for audio–visual but visual-tactile followed the same logic with tactile motion in place of auditory motion). Each experiment comprised three conditions: two unimodal blocks (one per modality) and an interleaved condition in which trials of the two modalities were randomly mixed. Each modality was assigned to its own range of speeds, with the two ranges partially overlapping. The common speeds are shaded grey. Colors denote modality throughout (green for vision; grey for visual-tactile). **(D)** Schematic of the central tendency effect and the regression index. Reproduced speed is plotted against actual speed; the dashed line shows veridical performance (slope = 1). Because estimates regress toward the mean of the presented range, slow speeds are overestimated and fast speed underestimated, so the fitted line is shallower than the veridical performance. We quantify this as the regression index RI = 1-slope, with the RI = 0 indicating veridical reproduction and RI = 1 complete regression to the range mean.

Within the Bayesian framework (Knill & Pouget, 2004; Petzschner et al., 2015), central tendency reflects the optimal combination of current sensory evidence with the prior probability distribution learned from environmental regularities (Cicchini et al., 2012; Jazayeri & Shadlen, 2010). When the uncertainty in sensory input is higher, the prior is weighted more and perceived features regress more towards the mean, resulting in greater central tendency. A key question is how central and general these priors are; does a single prior operate globally across sensory modalities or does each modality maintain its own prior? We recently addressed this by interleaving auditory and visual stimuli and found that perception maintained sensory-specific priors and did so for both spatial and temporal tasks (Tonelli, Phan, et al., 2025).

Building on this finding, the current experiment examines whether central tendency can be used as a behavioral probe to test whether two modalities encode a feature through a sensory-specific or supramodal process. To do so, we assign each modality its own range of stimulus values and interleave the presented modality. If priors are modality specific, the modalities should exhibit different central-tendency biases that resemble their own unimodally measured biases. If, on the other hand, the prior is supramodal, interleaving the two different stimulus ranges should produce a central-tendency bias reflecting the pooled set of speeds. Observing whether interleaved responses converge or remain separate therefore reveals whether the priors are supramodal or modality-specific.

The stimulus variable to be manipulated in this experiment is speed. Speed perception is critical to the safe and efficient navigation of the world and plays a key role in collision avoidance, sensorimotor coordination and interception. Crucially, it can be encoded by three different sensory systems (vision, audition, and somatosensation) and has been studied extensively in multisensory contexts (Gleiss & Kayser, 2014; Soto-Faraco et al., 2003; Tomassini et al., 2011; Tonelli, Burr, et al., 2025; Alais & Burr, 2004; Hanse, et al., 2019; Bensmaïa, et al., 2006). Speed perception is therefore a perfect candidate to test the validity of behavioral central tendency in probing feature processing and will allow for a comparison of visual-tactile and audio-visual conditions.

Although both vision and audition convey motion, they rely on very different neural mechanisms. In the visual system, motion is encoded by direction- and speed-selective neurons in primary visual cortex (Perrone, 2006; Priebe et al., 2006) and subsequently areas MT/MST, giving a precise, retinotopically organized representation (Burr & Thompson, 2011; Perrone & Thiele, 2001). The auditory system, by contrast, lacks dedicated motion detectors in early processing and instead speed is inferred based on spectral (i.e., frequency) and binaural cues (i.e., interaural difference) (Carlile & Leung, 2016; Freeman et al., 2014). Behaviorally, audio-visual motion is dominated by vision and shows little sign of optimal integration (Bentvelzen et al., 2009; Soto-Faraco et al., 2004; Tonelli, Burr, et al., 2025). Neuroimaging results are consistent with these findings, showing minimal cross-activation of visual motion area MT by auditory motion (Lewis, 2000; Rezk et al., 2020; Van der Stoep & Alais, 2020). Together, this suggests processing of visual and auditory motion is largely segregated and hence that modality-specific priors would be expected when interleaving auditory and visual motions. Vision and touch, in contrast, appear to share common processing. Several lines of evidence support this: adaptation in one modality produces corresponding aftereffects in the other (Konkle et al., 2009; Konkle & Moore, 2009), cross-sensory facilitation occurs between visual and tactile motion (Gori et al., 2011), and disrupting V5/hMT+, the region responsible for visual motion perception, impairs tactile motion discrimination (Amemiya et al., 2017) and tactile speed perception (Basso et al., 2012). In addition, both types of motion evoke common occipito-temporal activation (Haggard, 2017; Pei et al., 2011; Pei & Bensmaia, 2014). These overlaps are strong predictors of a shared motion processing and thus a single, supramodal prior for speed perception is expected for interleaved visual and tactile motion.

We will test these predictions (independent priors for vision and audition, a shared prior for vision and touch) in separate speed perception experiments employing the same behavioral paradigm in which each modality has its own range of speeds and unimodal priors are compared against the prior produced by interleaving the sensory modalities. To foreshadow the results, the interleaved condition produced a shared prior for visual-tactile speed but separate priors for audio-visual speed.

## Results

In each experiment, participants reproduced the speed of a moving stimulus, presented in one of two modalities (Figure 1). Each modality was given its own range of speeds but were partially overlapping. Observers completed two unimodal blocks (one per modality) and a third interleaved block in which the two modalities were randomly interleaved. If each modality maintains a separate prior, the speed biases should remain tied to their own range even when interleaved. If the two modalities share a supramodal prior, a single bias should emerge as the two speed ranges will converge on a common mean. A key point is what happens to the common speeds in the overlapping portion of the speed range. If there are separate priors, these common speeds should be reproduced differently in each modality as they will be biased towards different means.

Figure 2 shows the central tendency data for all unimodal conditions. Reliable central tendency biases were present in every modality (visual, auditory and tactile), and the bias for vision was present in both the audio-visual and visual-tactile conditions. The central tendency bias is due to the reproduced speeds regressing towards the mean of the speed range and can be quantified by a regression index, as shown in Figure 1D. The regression indices (RIs) for all conditions shown in Figure 2 are significantly above zero (*p*’s < 0.001).

**Figure 2.**
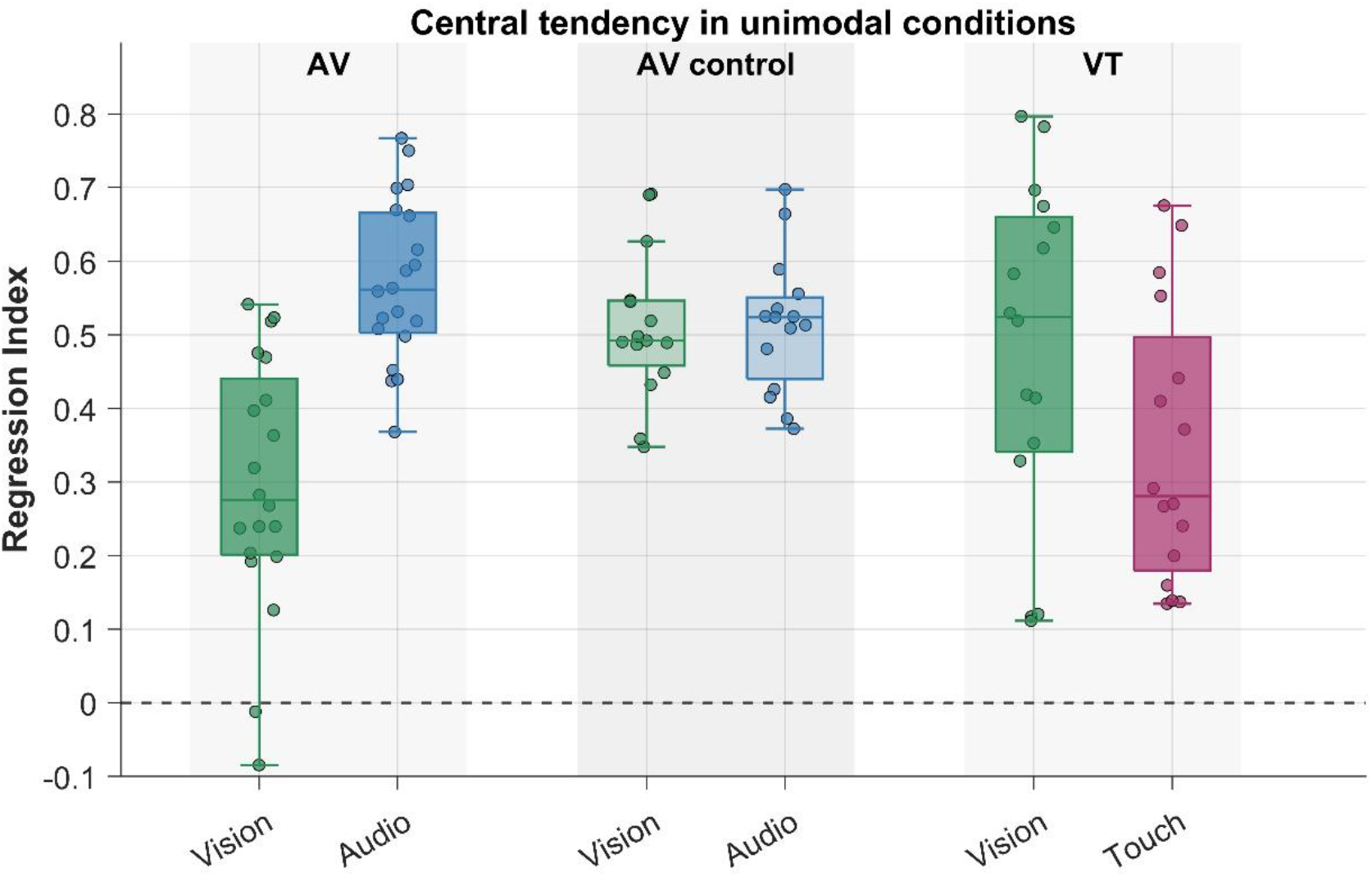
Central tendency in unimodal conditions. Regression index (RI = 1 − slope of reproduced against physical speed) for each modality in the three experiments (AV, n = 20; AV control, n = 15; VT, n = 16). The horizontal dashed line shows RI = 0 and denotes veridical reproduction, whereas RI values greater than 0 indicate regression towards the mean of the speed range (i.e., the central tendency effect) with a value of 1 corresponding to complete regression. Box-and-whisker plots show the median and interquartile range (box) and the overall range (whiskers) with outliers excluded, and dots show each individual’s RI value. Colors denote modality (green, vision; blue, audition; magenta, touch). The lighter colors in the central group represent the precision-control experiment in which visual precision was reduced to equate visual and auditory RIs. Central tendency was reliably present in every modality and experiment (all RI’s are significantly greater than 0, all *p*’s < 0.001).

The results are also consistent with a reliability-based Bayesian account because the degree of central tendency was stronger when reliability was low. For each participant and modality we estimated sensory uncertainty from the unimodal conditions as the variability of residual of the linear fit between the reproduced speed and the actual speed, corrected for the compression that central tendency itself introduces (see Data Analysis for more details). On this measure, audition was markedly noisier then vision in the audio-visual experiment (uncertainty fold-gap = 13.4), whereas vision and touch were more closely matched (uncertainty fold-gap = 2.5). In the audio-visual experiment, regression to the mean in the unimodal conditions was greater for the less reliable modality (i.e., audition) (RIaudio > RI_vision_, *t*_19_ = 7.7, *p* < .001, Cohen’s *d* = 1.72). Similarly, in the visual-tactile experiment, unimodal regression to the mean was greater for touch: RI_touch_ > RI_vision_, *t*_15_ = 3.08, *p* < .01, Cohen’s *d* = 0.77). This pattern of results suggests a greater reliance on the prior (the mean speed) when sensory input was less reliable, consistent with a Bayesian account.

The reliability of the speed ratings differed among the unimodal conditions, and we quantified this by calculating the uncertainty fold-gap: the ratio of the larger to the smaller of the two uncertainties defined above. In the audio-visual condition, the mismatch was large because auditory speed estimates were far more dispersed than visual estimates. The median value of the uncertainty fold-gap was13.4 (i.e., audition was roughly thirteen times less reliable than vision). In the visual-tactile condition, the two modalities were more closely matched in sensory uncertainty (median fold-gap = 2.5). The very large difference in relative reliability could be problematic because, even though Bayesian cue combination holds that a modality should be pulled toward a shared prior by an amount inversely proportional to its own reliability, it is possible that this principle could fail for two sensory modalities of extremely different reliabilities. With this in mind, we conducted a precision-control experiment (see Methods) designed to match the visual and auditory reliabilities by adding noise to the visual stimulus. The box-and-whisker plots shown in the centre of Figure 2 show that the control experiment was successful in producing very similar reliabilities for audition and vision as the overall range and the interquartile range were very similar in each condition. Moreover, once approximately matched in reliability, the two conditions produced equivalent degrees of central tendency bias and the difference between their RI values was not significant.

To probe the shared priors, we analyzed the common speeds that were physically identical in each pair of modalities. When vision and audition were interleaved (Figure 3, top row), the speed reproductions depended on modality (visual vs auditory) and condition (unimodal vs interleaved bimodal) in a significant interaction (F_(1,4864)_ = 22.9, p < .001), which we unpacked with four planned comparisons: the between-modality difference and the within-modality change between unimodal/interleaved condition. Speed reproductions differed significantly between the two modalities: the speed-bar ratings differed by -279 pixels in the unimodal blocks (p < .001, dz = 2.27) and, although this separation was significantly reduced when the modalities were interleaved, it remained large at -186 pixels (p < .001, dz = 1.48). A Bayes factor confirmed that this interleaved between-modality difference was itself far from zero (BF10 = 7.8 × 10^3^ for a non-zero interleaved difference). The reduction in modality difference was driven by only modest within-modality shifts: visual responses moved reliably in the direction that narrows the gap toward audition (p = .005, dz = 0.74, BF10 = 12), whereas audition showed no clear shift (dz = 0.42, BF10 ≈ 1, inconclusive). Thus, the modality difference in the audio-visual pairing was reduced but preserved by interleaving, and the small shift that occurred was confined to one modality. Overall, this reveals a partial and asymmetric convergence rather than the speed estimates in each modality shifting towards a common value.

**Figure 3.**
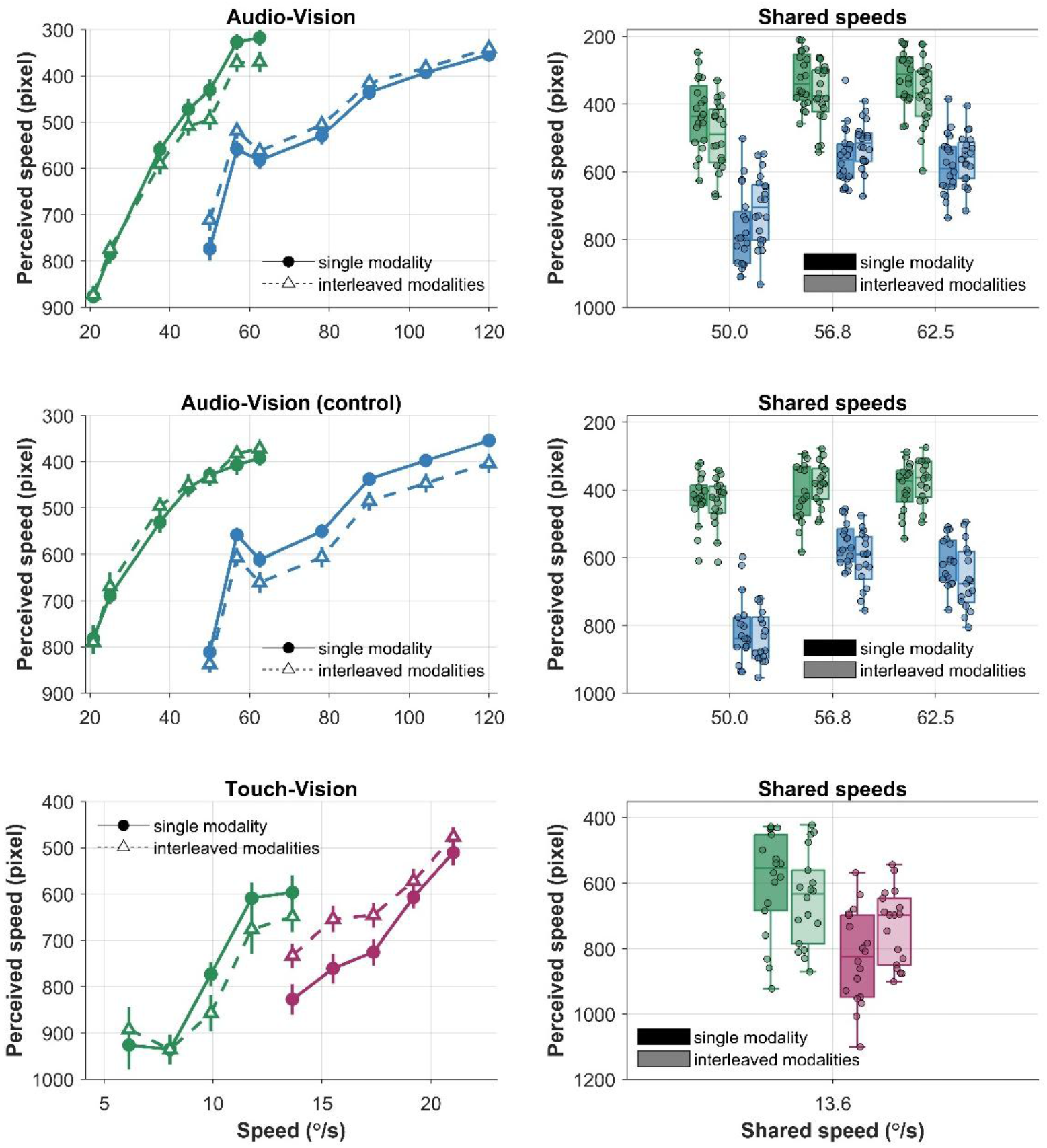
Reproduced speed in the unimodal and interleaved conditions. Rows correspond to the three experiments (top, audio–vision; middle, audio–vision precision control; bottom, tactile–vision). *Left column:* mean reproduced speed, expressed as bar position in pixels (note the reversed y-axis, with faster speeds toward the top), as a function of physical stimulus speed, for each modality in the unimodal (solid lines, filled circles) and interleaved (dashed lines, open triangles) conditions. Error bars are ±1 SEM. Each modality was assigned its own speed range but partially overlapped with common speeds. Colors denote modality (green, vision; blue, audition; magenta, tactile). *Right column:* distribution across participants of reproduced speed at the shared speeds (50, 56.8 and 62.5 °/s for audio–vision; 13.6 °/s for tactile–vision), for each modality in the unimodal (darker fill) and interleaved (lighter fill) conditions. Boxes show the median and interquartile range, whiskers the range excluding outliers, and dots individual participants. For the shared speeds, the between-modality difference is largely preserved from the unimodal to the interleaved block in the audio–visual condition (and, if anything, is larger in the precision-matched audio– visual control condition) whereas in tactile–vision the difference collapses and the two modalities converge on a common value.

Conversely, the visual-tactile speed data (Figure 3, bottom row) behaved in the opposite way and showed a clear supramodal convergence. The interaction between modality (visual vs tactile) and condition (unimodal vs interleaved bimodal) was significant (F_(1,1256)_ = 52.3, p < .001). Speed reproductions were very different between the two modalities when unimodally presented (−302 pixels, p < .001, dz = 1.47, BF10 = 807) but this separation largely collapsed when they were interleaved, shrinking by 72% to -86 pixels. Here, the evidence was equivocal as to whether the between-modality difference remained (dz = 0.52, BF10 = 1.4, inconclusive), and across participants the interleaved difference was no longer significant (p = .10). Unlike audio–vision, both modalities moved substantially from their own baseline to converge in the middle, with strong evidence for each shift: vision by -88 pixels (dz = 0.84, BF10 = 11) and tactile by +127 pixels (dz = 0.98, BF10 = 29). As a reliability-weighted shared prior predicts, the less reliable modality (touch) showed a greater degree of convergence. This pattern of data – a between-modality difference that largely collapses when interleaved and both modalities shifting towards each other to a common estimate – indicates that visual-tactile speed forms a shared prior. There is, though, a small, statistically inconclusive residual that the convergence was substantial but not complete.

Because speed reproduction for audition was so much less reliable than for vision, this large reliability ratio may have been what prevented us from finding evidence of a supramodal audio-visual prior. To be sure that the failure to find a supramodal prior was a genuine result, we repeated the audio-visual experiment after making vision less precise so that the audio-visual reliability ratio would be similar to that of the visual-tactile pairing, and which did reveal a supramodal prior. The visual-noise manipulation succeeded in reducing visual reliability: mean visual uncertainty increased from 142.34 (SD = 68.09) in the main experiment to 396.47 (SD = 238.87) in the control (Mann–Whitney, p < .001), and the audio-visual reliability ratio shrank from a 13.4-fold to a 3.2-fold ratio — statistically equivalent to the 2.5-fold ratio of vision–touch (TOST on the log ratio, factor-of-two margin, p = .021). If it were the large reliability ratio that prevented the supramodal audio-visual prior, the reliability-matched control should now behave like the visual-tactile condition and converge on a supramodal prior. In fact, it did the opposite (Figure 3, central plot). The two modalities differed by −248 pixels when measured separately in the unimodal conditions (p < .001, dz = 2.56) and by −302 pixels when interleaved (p < .001, dz = 3.41). The interaction between modality (audition vs vision) and condition (unimodal vs interleaved bimodal) was significant (F(1,4190) = 20.9, p < .001), and a Bayes factor confirmed that the interleaved between-modality difference was far from zero (BF10 = 3.8 × 106 for a non-zero interleaved difference). Vision did not shift (+19 pixels, p = .22, dz = 0.31, BF01 = 2.0, evidence for no shift) but audition moved away from vision (−35 pixels, p = .002, dz = 0.65). Thus, even when matched for reliability, we found no evidence for a supramodal prior.

To place all three datasets on a single comparable scale, we computed for each participant a between-modality *convergence index*: in essence, how much of the unimodal separation between the two modalities is lost once they are interleaved. The index ranges from −1 to +1, with zero meaning the separation was unchanged, positive values meaning the modalities converged (1 = complete convergence), and negative values meaning the modalities separate further in the interleaved block (see Methods for details). The index provides a descriptive summary of convergence, not an estimate of how strongly a shared prior is weighted. The index captured the graded pattern (Fig. 4): visual-tactile converged (median C = +0.45, 95% CI [+0.27, +0.58]), the audio–visual main experiment showed reliable but partial convergence (+0.17 [+0.09, +0.36]), and the audio–visual control showed increased separation (−0.13 [−0.17, −0.02]). The two audio-visual datasets bracket the comparison: the reliability-matched control was clearly more separated than visual-tactile (Wilcoxon p = .0001), while the main audio-visual experiment, with its partial convergence, sat between the two and did not differ significantly from visual-tactile on this index (p = .18). The clearest contrast is therefore with the control, where audio-visual and visual-tactile have equivalent reliability ratios yet diverge in how their behavior when interleaved, with only visual-tactile showing clear evidence of supramodal priors.

## Discussion

Using central tendency as a behavioral probe, we asked whether speed priors are shared across modalities or kept modality-specific. We did so allocating to each modality a different but partially overlapping range of speeds and comparing unimodal speed reproduction with bimodally interleaved reproduction. When vision was combined with touch, interleaving caused the speeds common to the two modalities to converge to a large extent. In fact, in the interleaved block, the difference between the modalities decreased by 72%, and both modalities shifted towards each other, with touch shifting more in a manner expected of a reliability-weighted shared prior. When vision was paired with audition, the difference between modalities instead persisted and audition, the less reliable modality, failed to converge. Interestingly, this audio–visual segregation was not a result of the large reliability gap between the senses. It was still observed when visual precision was reduced to make the audio-visual reliability ratio similar to that of the vision–tactile range which did produce supramodal convergence. Indeed, reducing visual precision made the audio– visual estimates more segregated rather than more convergent, contrary to supramodal account. Therefore, whether central-tendency priors are shared across modalities appears to be determined by which senses are paired rather than the relative reliability of those senses.

These patterns fit the idea that whether two modalities recruit a shared or separate prior(s) is dependent on the compatibility of their underlying processing. Visual and tactile motion both encode spatial and temporal properties at a primary level, and both draw on overlapping processing with shared occipito-temporal activation and causal V5/hMT+ involvement in tactile motion (Amemiya et al., 2017; Basso et al., 2012; Gori et al., 2011; Pei et al., 2011). These commonalities may be key elements which permit a supramodal prior to be calculated. In contrast, visual and auditory motion rely on largely distinct codes anchored in different reference frames (Carlile & Leung, 2016; Rezk et al., 2020; Van der Stoep & Alais, 2020) and audition lacks the primary spatial encoding that are present in vision and audition. Audition therefore computes motion at a higher-level than the other senses as it is an inference based on a collection of cues. Moreover, calculating auditory motion is a sluggish process, and it lags considerably behind visual motion processing (Tonelli, Burr, et al., 2025). These differences may be enough to prevent the pooling of auditory and visual speed signals into a supramodal prior. Our experimental design does not isolate this mechanism as the two pairings (i.e., audio-visual and visual-tactile) differed in their speed ranges. Future work could examine this further and employ neurophysiological measures.

Beyond any specific result, the study establishes central tendency as an implicit, non-invasive read-out of prior influence, one that requires no explicit judgements and can be applied to any feature that is shared across modalities. In quantifying how far estimates regress toward the supramodal mean across contexts, it can be inferred whether observers rely on shared or separate priors without neural data. That said, this study has some limitations. For example, the dissociation between cross-modal pairs confounds sensory modality with stimulus speed, an element that should be kept in mind when interpreting the current findings. Regardless, we did rule out a reliability-based account using a precision-matched control experiment. Another limitation is that the findings are behavioral and that the interpretation based on processing compatibility is a hypothesis supported by previous neuroimaging studies rather than a claim demonstrated in this study. However, this hypothesis is entirely testable.

In summary, whether speed priors are shared across the senses depends on which senses are paired. What determines which sensory pairs come to share a prior is the question these results open. We conjecture that a fundamental featural compatibility is necessary for two senses to pool their stimulus history into a supramodal prior.

## Methods

### Participants

For the vision–audition experiment, twenty-one participants were tested (age 21.14 years, SD = 3.57) and twenty analyzed after exclusions; for its precision-control variant, seventeen were tested and fifteen analyzed (age 19.94 years, SD = 1.297); for the vision–touch experiment, eighteen were tested (age 21.17 years, SD = 4.76) and sixteen analyzed. All had normal or corrected-to-normal vision and normal self-reported hearing or tactile sensitivity and gave written informed consent (HREC 2021/048). Participants were excluded if flagged as outliers (> 3 SD from the mean) in either modality’s baseline sensory uncertainty, or if the slope of their reproduced-speed versus actual-speed function was negative (i.e., a non-monotonic mapping, with faster stimuli reproduced as slower).

### Stimuli and apparatus

Stimuli were generated in MATLAB (R2017a for vision–audition; R2022b for vision–touch) with Psychtoolbox (Brainard, 1997) in a dark, sound-attenuated room. In the vision–audition experiment, visual stimuli were front-projected on a ∼55° white screen using a PROPixx projector at 120 Hz (resolution 1920 x 1080 pixels); a ∼4° cosine-edged circle at 50% contrast moved horizontally. Auditory stimuli were white noise played over eleven speakers (Visaton F8SC) spaced ∼5° apart behind the screen, with smooth apparent motion produced by convolving the noise with speaker-specific cosine windows. Visual speeds averaged ∼42°/s (20.8–62.5°/s) and auditory speeds ∼80°/s (50–120°/s). These speed ranges had three overalpping values (50, 56, 62.5°/s). In the visual precision control condition, the visual speeds were identical but visual precision was reduced by adding a spatially filtered visual noise background (0.14–0.43 cycles/deg, temporally white, 30% contrast) alpha-blended (0.25) onto the target using the *BlendFunction* in Psychtoolbox. The addition of visual noise reduced the reliability of the visual condition so that the large modality difference in reliability in the audio-visual condition was reduced to match that of the visual-tactile condition. In the vision–touch experiment, visual and tactile motion were delivered by two custom rotating wheels (10.5 cm, 3D-printed PLA, sinusoidal ridges 30 mm wide, 2.5 mm deep, 3.5 mm spacing), driven by high-precision stepper motors (1.8°/step) via an Arduino interface. The tactile wheel was ∼45 cm and the visual wheel ∼35 cm from the head. Visual speeds averaged ∼9.8°/s (6.1–13.6°/s) and tactile surface speeds ∼17.3°/s (13.6–21°/s), with one shared value (13.6°/s).

### Procedure

Participants estimated the speed of each stimulus by adjusting a black square (1°) along a vertical bar (∼24°) using the keyboard’s arrow keys. The upper end of the speed bar indicated “fast” perceived speed and the lower end “slow” and the space bar was used to submit the response. There were eight practice trials with feedback preceding each experiment. Each experiment comprised three conditions in random order: two unimodal conditions (one per modality) and an interleaved condition in which the two modalities were randomly intermixed. Each modality had different speed ranges which ensured that their mean speeds (on which the unimodal priors should be centered) also differed. In the interleaved condition, if a supramodal prior emerges, it should be centered at an intermediate point between the unimodal means. A key design feature is that the speed ranges partially overlapped. These common speeds help distinguish between unimodal and supramodal priors as they may exhibit a bias towards the unimodal mean or the supramodal mean. Indeed, based on our predictions, the same visual speed should be biased to the supramodal mean in the audio-tactile case and to the visual mean in the audio-visual case. Each speed was repeated 21 times (vision-audition: 147 trials per unimodal block, 294 interleaved; vision-touch: 105 trials per unimodal block and 210 interleaved).

### Data analysis

Trials with reaction times below 200 ms or above 3 SD over the condition median were removed. Analyses used R (lme4, lmerTest, emmeans, BayesFactor), with Satterthwaite degrees of freedom. For the unimodal conditions, responses on the speed bar were rescaled to speed per modality (°/s) to obtain the regression index (RI = 1 − slope; 0 = veridical, 1 = full regression to the mean). We tested RI against zero (one-sample t-tests) to confirm that central tendency was present and compared RI between the two modalities of each experiment (paired t-tests) to relate its strength to relative reliability. Moreover, for each participant and modality we estimated sensory uncertainty from the unimodal conditions, following Aston et al. (Aston et al., 2021). For each participant we fitted a linear regression of the reproduced speed on the actual speed and computed the variance of the residuals about that fit. Because central tendency compresses responses toward the range mean, this residual variance is itself compressed by the same factor, so it was corrected by dividing it by the squared slope of the fit, σ^2^_sensory = var(residual)/slope^2^. This yields the trial-to-trial variability of a modality’s estimates once the compression due to central tendency is removed; larger values indicate a noisier, less reliable modality. To capture the balance of uncertainty between the two modalities within each experiment, we then summarized each participant with an uncertainty fold-gap: the ratio of the larger to the smaller of the two modalities’ uncertainties (1 = equally reliable; larger values = one modality much noisier than the other).

In the interleaved condition, the raw speed bar responses (in pixels) were analyzed (they were not rescaled) because the mapping between bar position and physical speed is precisely what distinguishes the competing hypotheses; stimulus speed entered only as a predictor. For the common speeds shared among the modalities (three in the audio-visual conditions and one in the visual-tactile condition), we fitted linear mixed models of the form: pixel ∼ modality × session × speed + (1 + session | subject). Speed was omitted for the visual-tactile condition as there was only a single shared speed, reducing the random structure to an intercept when a model was singular. The modality × session interaction tests whether the modality difference changes from baseline to interleaved. From each model, we estimated a single family of four pre-specified contrasts: the modality difference within each session, and the baseline-versus-interleaved shift within each modality. We applied Bonferroni correction to the p-values across the family. For every contrast we report a standardized effect size (Cohen’s dz on the per-participant values) as well as a Bayes factor (ttestBF on the per-participant values, medium Cauchy prior r = 0.707; robustness checked across r = 0.5–1.414) so that null outcomes, for example, a modality that does not shift, are quantified as evidence rather than being treated as mere non-significance. To summarize convergence per participant on a bounded scale we used:

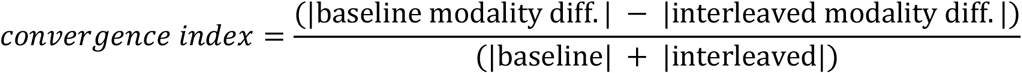

The convergence index lies in the range [−1, +1] where: 0 = separation retained, 1 = complete convergence, < 0 = increased separation. The convergence indices for audio-visual and visual-tactile conditions were compared with Wilcoxon rank-sum tests. The reliability imbalance in each sensory pairing was quantified as the per-participant uncertainty fold-gap (ratio of the larger to the smaller unimodal sensory variance). To verify that the manipulation worked in the audio-visual control experiment, we compared visual sensory uncertainty of the main and control audio-visual experiment (Mann–Whitney), confirming that adding visual noise did reduce visual reliability; we then tested whether the resulting audio-visual uncertainty ratio was equivalent to the visual-tactile one with a TOST (two one-sided t-tests) on the log fold-gap (factor-of-two margin). As an assumption-light complement, paired non-parametric tests on per-participant means confirmed whether the modality difference survived in the interleaved block.

## Acknowledgement

We thank Yixi Wen for assistance with part of the data collection.

## Fundings

This work is funded by the European Union Horizon Europe research and innovation programme under the Marie Skłodowska-Curie-2021-PF-01 (FLEX-U - g.a. No. 101064748). The views and opinions expressed are those of the author(s) only and do not necessarily reflect those of the European Union. Neither the European Union nor the granting authority can be held responsible for them.

## Notes

### Competing Interest Statement

The authors have declared no competing interest.

